# Deep Neural Networks for Genomic Prediction Do Not Estimate Marker Effects

**DOI:** 10.1101/2021.05.20.445038

**Authors:** Jordan Ubbens, Isobel Parkin, Christina Eynck, Ian Stavness, Andrew Sharpe

## Abstract

Genomic prediction is a promising technology for advancing both plant and animal breeding, with many different prediction models evaluated in the literature. It has been suggested that the ability of powerful nonlinear models such as deep neural networks to capture complex epistatic effects between markers offers advantages for genomic prediction. However, these methods tend not to outperform classical linear methods, leaving it an open question why this capacity to model nonlinear effects does not seem to result in better predictive capability. In this work, we propose the theory that, due to a principle called shortcut learning, deep neural networks tend to base their predictions on overall genetic relatedness, rather than on the effects of particular markers, such as epistatic effects. Using several datasets of crop plants (lentil, wheat, and *Brassica carinata*), we demonstrate the network’s indifference to the values of the markers by showing that the same network, provided with only the locations of matches between markers for two individuals, is able to perform prediction to the same level of accuracy.

## Introduction

Genomic Prediction (GP) is the practice of predicting trait values, such as yield, for members of a population using marker data. In the breeding of crop plants, GP has been shown to be an effective way to speed the rate of improvement for agronomically relevant traits (1–3). GP has become increasingly accessible in recent times due to rapid improvements in genotyping technologies using next generation sequencing at a much reduced cost, and has been integrated into many breeding programs around the world (4).

The literature on GP in plants and animals is mature, with linear additive models of genetic effects being the most popular in practice. However, recent years have seen an increasing interest in using machine learning methods, including deep learning, for the GP task. Although it is widely assumed that deep neural networks will learn to model complex nonlinear epistatic effects (5), they rarely perform better than simple linear models such as the *Genomic Best Linear Unbiased Predictor* (GBLUP) (6). This implies that deep neural networks either do not estimate the effects of specific alleles, or do not benefit from doing so.

In this work, we explore the gap between the capacity and the real-world performance of deep learning methods for GP by controlling what information the network is allowed to access. We develop a novel technique for training a deep neural network which performs prediction based on the locations of matches between markers for pairs of individuals, without giving the network access to the marker data directly. Under the reasonable assumption that a deep neural network estimates marker effects, this training strategy is expected to perform poorly as the network has no information about which alleles are present in any individual. However, through experiments with different plant species and population structures, we show that this prediction strategy, based solely on genetic relatedness, performs just as well as allowing the network to access the values of the markers. The equivalence in predictive performance for deep neural networks with or without access to the marker data implies that these networks tend to rely on estimating overall genetic relatedness, not the effects of specific markers.

## Background

### Kernel Methods

The term *kernel method* refers to a class of models which rely on a fixed or learnable kernel function *K* which relates a notion of “similarity” between its two inputs. Kernel machines are a common example of kernel methods in machine leaning. In the simplest terms, kernel machines are models in the form

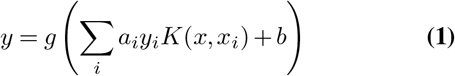

where *x* and *y* are the input and output of the model respectively, (*x*_*i*_, *y*_*i*_), *i* ∈ { 0 … *N* are the *N* samples of the training set (the data vectors and the labels), *g* is a an optional transfer function, and *a_i_, i* ∈ {0 *… N*} and *b* are learnable weight and bias terms. In this way, the output of the kernel machine for an input is a function of the weighted similarity between it and the training samples, as indicated by the kernel function *K*. Kernel functions have classically been chosen based on expert knowledge of the data or the problem space, for example, using Gaussian or polynomial kernels.

In GP, the *Genomic Best Linear Unbiased Predictor*, or *GBLUP*, is a common estimator of the form

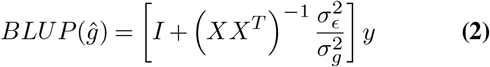

where *X* is the (mean-centered and normalized) matrix of markers, and *σ_E_* and *σ_g_* are the standard deviation of errors and genetic effects, respectively (7). Here, the covariance matrix *XX^T^* represents a simple linear kernel, capturing linear additive effects among markers. Nonlinear kernels can also be used in the estimation of genetic relatedness. *Reproducing Kernel Hilbert Spaces* (RKHS) regression with a Gaussian kernel has seen some success in the literature, although linear kernels still remain the most popular (4).

### Neural Networks and Deep Learning

Nonlinear models such as neural networks have the theoretical capability to capture arbitrarily complex epistatic effects in whole-genome regression. For example, multiple markers may have negligible effects individually, but significant effects when considered in combination. The ability to model these interactions gives nonlinear models a potential representational advantage over linear models for prediction. Several authors have suggested that this advantage makes deep neural networks promising candidates for performing GP (5, 8, 9).

Machine learning methods for GP, including deep learning methods, have been explored in the literature. The results have been mixed, with the performance of deep learning varying between datasets. One of the most commonly used neural network architectures is the Convolutional Neural Network (CNN), which uses layers of filter banks to capture local patterns in the data. Despite their representational power and their success in other subfields of biology, CNNs have been shown to perform similarly to classical linear models in some cases, although somewhat under-performing them in others (6). One extensive survey of GP methods for various traits in six different plant species showed that CNNs performed the worst among all tested models on average (10). Another study in soybean showed similar performance between deep learning and linear models when missing SNPs were imputed (11). Results for blueberry and strawberry again saw no advantage for CNNs over conventional methods for most traits (12).

The similarity in performance between powerful, high-capacity deep learning models and standard linear models suggests that either the genetic architectures of all the examined traits are strongly linear additive in nature, or that deep neural networks are not estimating nonlinear effects between markers despite their ability to do so.

## Methods and Materials

### Obscuring Marker Data from the Network

Due to the black-box nature of deep neural networks, it is difficult to determine precisely how the network arrives at a prediction (13). Therefore, the most reliable way to determine whether the network benefits from estimating the effects of particular alleles is to completely disallow the network from accessing the values of the markers themselves.

To investigate whether deep neural networks estimate phenotype based on estimating marker effects, we propose a novel training method which obscures the marker data from the network. A schematic of the proposed method is shown in Figure 1. Comparing to the kernel machine formulated in Equation 1, the proposed model can be represented as

**Fig. 1.**
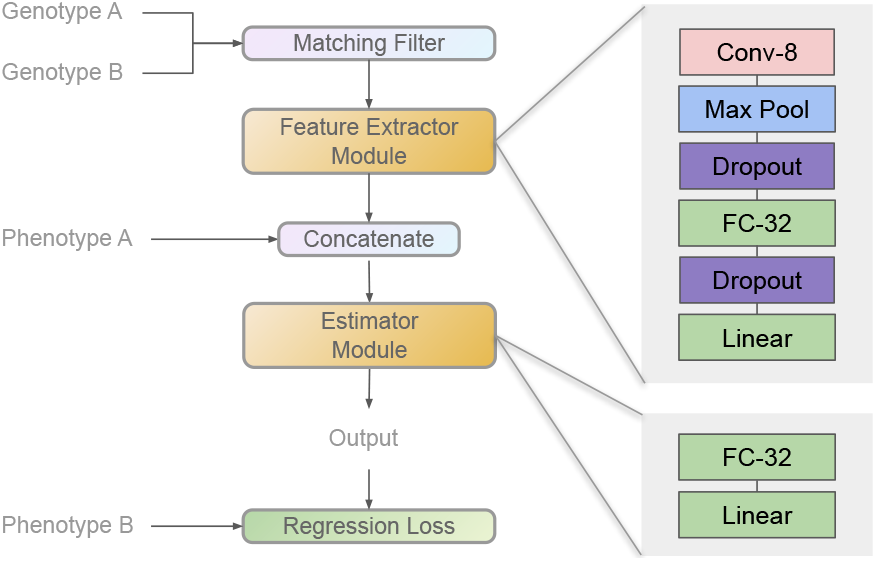
Overview of the proposed method for training a deep neural network without giving the network access to the marker data. Inset, the DeepGS architecture (5) is shown as the feature extractor module, and a two-layer fully-connected network is shown as the estimator module.

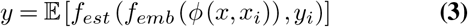

where *ø* is the matching filter operation

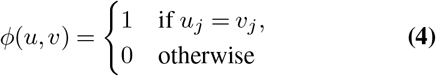

for each marker *j*, *f_emb_* is the *feature extractor module* and *f_est_* is the *estimator module*. The matching filter only outputs one when the encodings match exactly, so comparing heterozygous and homozygous individuals will produce a zero. Here, the subscripted (*x_i_, y_i_*) is a member of the training population, called the *reference sample*, and (*x, y*) is the sample *x* for which the phenotype *y* is being predicted, called the *query sample*. In a manner similar to kernel machines, the final point estimate produced for a query sample is an aggregate of predictions made over all members of the training population.

During the training phase, the feature extractor module and the estimator module are trained in an end-to-end fashion using random pairs of samples from the training population. The training samples are retained for inference time, during which a forward pass is performed on each query sample using each member of the training population as the reference sample. Because the output is a distribution over predictions, the point estimate is given by the mean of this output distribution. Various other aggregation methods such as ensemble learning and mixture-of-experts models (14) were trialed, with no consistent improvement over using the simple mean. In practical terms, any nonlinear models can be used for the feature extractor module and for the estimator module. Here, we use a simple two-layer feed-forward neural network for the estimator, and we use the DeepGS convolutional neural network architecture from the literature for the feature extractor module (5). DeepGS is a small convolutional architecture, using a single convolutional layer and two fully connected layers. It uses stochastic regularization in the form of Dropout (15). DeepGS was selected as it is the most commonly used example of a deep network built specifically for GP. We use DeepGS here as this allows us to make direct comparisons to the DeepGS architecture used in a whole-genome regression approach, as differences in performance are therefore not to due to differences in architecture or model capacity.

The proposed method is distinct from RKHS methods in the fact that Equation 3 is not a reproducing kernel *K* (it is not guaranteed that *K* : *X* × *X* → ℝ is positive semi-definite). It is also distinct from standard whole-genome regression methods, as the network never has access to the marker data. It is only provided with the locations of matches between two otherwise anonymous samples, and the specific alleles of both samples are unknown. This means that the network cannot reduce to existing whole-genome regression methods by regressing on the markers of the query sample while ignoring the reference sample.

### Models Evaluated

To test our hypothesis, we compare the proposed method, with DeepGS as the feature extactor, to DeepGS used in the typical fashion, performing whole-genome regression with direct access to the marker data. For completeness, we also compare these networks to two common linear models, GBLUP and LASSO, as well as two other machine learning methods: a fully-connected neural network (FC), and a fully-connected neural network with *L*_0_ regularization (16).

LASSO is a penalized linear model of the form

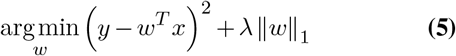

where *w* are the model parameters and *λ* is a tuneable constant which controls the strength of the regularization term. In practice, the *L*_1_ penalty on the weight vector tends to encourage sparsity in the weights, a desirable quality under the assumption that a low proportion of the total markers have influence on the trait. LASSO and a related model, the Elastic Net, performed the best (and better than the commonly used RR-BLUP) in a survey of penalized linear models for GP (17).

Fully-connected neural networks, also known in some GP literature as Multi-Layer Perceptrons (MLPs), are a common choice for machine learning applications in GP. We use a standard two-layer neural network. Batch normalization is used, as it potentially improves generalization performance and allows training to succeed over a higher range of learning rates.

Motivated by the principle of sparsity, we also investigate a fully-connected neural network equipped with *L*_0_ regularization (16). While *L*_1_ regularization penalizes the absolute magnitude of the weights, *L*_0_ regularization penalizes the proportion of non-zero weights, explicitly encouraging sparsity. Optimizing this regularization term using gradient descent methods is impossible, because the *L*_0_ norm is not differentiable. However, it can be approximated through a *hard concrete distribution* - a continuous relaxation of discrete random variables (18, 19). The result is differentiable feature selection, learned simultaneously with the model (20).

### Hyperparameter Selection

In order to fairly represent the performance of each method, care must be taken to tune them individually for optimum performance. In the machine learning literature, constants used in the model specification or during training, such as the learning rate or the size in units of a hidden layer, are referred to as *hyperparameters*. The process of selecting the values for hyperparameters which are likely to result in the best performance on the test data is known as *hyperparameter tuning*. This is done by holding out a portion of the training data, called the *validation set*. Performance on the validation set can be used as a proxy for performance on the unseen test data, allowing an automated search over values of the hyperparameters in either a random or a structured fashion. In this section we describe how hyperparameters were selected for each of the six methods. For the LASSO model, the regularization coefficient *λ* = 0.001 is used, as this value allows convergence in every dataset. Tuning *λ* resulted in slightly worse performance overall, and the bounds for tuning must be set carefully or values for *λ* may be selected which cause non-convergence when training the final model. For the FC and *L*_0_ models, the number of hidden units and the learning rate are determined via hyperparameter search as this consistently improved performance over predetermined constants. We use an exponentially decaying temperature schedule for the *L*_0_ model (20). For the DeepGS network, most hyperparameters are prescribed by the authors of the architecture. We use a learning rate of 1 × 10^−5^. Tuning the learning rate does not improve performance, and sometimes results in learning rates which are infeasible for training the final model.

For the proposed model, hyperparameters were predetermined as hyperparameter tuning proved to be both costly and the performance gains inconsistent. We use a value of *d* = 1 for the dimensionality of the embedding emitted by the feature extractor module. Smaller values of *d* tend to be better for small datasets, and so we find this to be the most general. The number of hidden units in the estimator module is set to 32, as tuning did not change performance significantly.

In addition to hyperparameter tuning, the validation set can also be used for *early stopping*. Some models may see a decrease in error on the validation set until a cert in number of training epochs, and increasing error thereafter. Halting training at this inflection point can prevent the model from overfitting the training data. However, in all experiments presented here, we found that it was universally better to use all of the training data to train the model, instead of reserving a portion as a validation set to monitor for early stopping. This is consistent with previous reports that the size of the training population is one of the most important factors in predictive performance (9). For all models, we add the validation set back in to the training data after hyperparameter tuning (when applied) and use the entirety to train the final model. Each model was trained until the training error had ceased to decrease.

### Datasets

In order to evaluate the difference between deep neural networks trained with and without access to the raw marker values, we use four datasets of food crops, representing a variety of species and sample sizes. Sources for the data are provided in the final section.

The first two datasets we use for evaluation are published datasets of lentil (*Lens culinaris*) (21). The authors of this study examined GP performance within-population, across-populations, and across-environments using a variety of models. The authors published a lentil diversity panel (LDP) made up of 320 individuals, as well as two biparental populations, termed LR-01 and LR-11. To evaluate the largest and most genetically diverse population possible, we use the combined LR-01 and LR-11 dataset from the paper, constituting a total of 230 individuals. This data is an amalgamation of two different biparental populations, and the authors of the data reported lower predictive performance on average using the two populations combined as opposed to using each individually. This makes an interesting case study, given the unusual population structure in this challenging combined population. The 39k SNPs sampled via exome capture provided by the authors are used.

We also use a population of 2,536 *Brassica carinata* plants. This population is a nested association mapping (NAM) population consisting of recombinant inbred line (RIL) progeny from 50 crosses of different *B. carinata* lines with a common reference line. This dataset includes 17k SNPs.

For the final crop dataset, we use a published dataset of 2,403 landrace accessions of Iranian wheat (*Triticum aestivum*) (22). The individuals were phenotyped for a variety of agronomically relevant traits and the authors of the dataset used these for a GP study with standard models. This is the dataset previously reported in the DeepGS paper, and so we include it in order to be consistent with the analysis from that publication (5). The dataset includes a set of 33k SNPs obtained via genotyping by sequencing (GBS).

Each dataset was split into five folds, and each model was trained five times using each fold as the hold-out set. Hyperparameter tuning was performed within each split when applicable, using one of the four training folds as the validation set.

## Results

Prediction results for the lentil diversity panel, lentil biparental population, carinata, and wheat datasets are shown in Figure 2, Figure 3, Figure 4, and Figure 5, respectively. As is usual in GP, we report Pearson’s *r* between the predicted and actual phenotypes as the measure of predictive performance. A property of *r* which should be mentioned is the distribution of *r* values for two independent random variables. For samples drawn from two independent Gaussian distributions, the probability density for *r* is given by

**Fig. 2.**
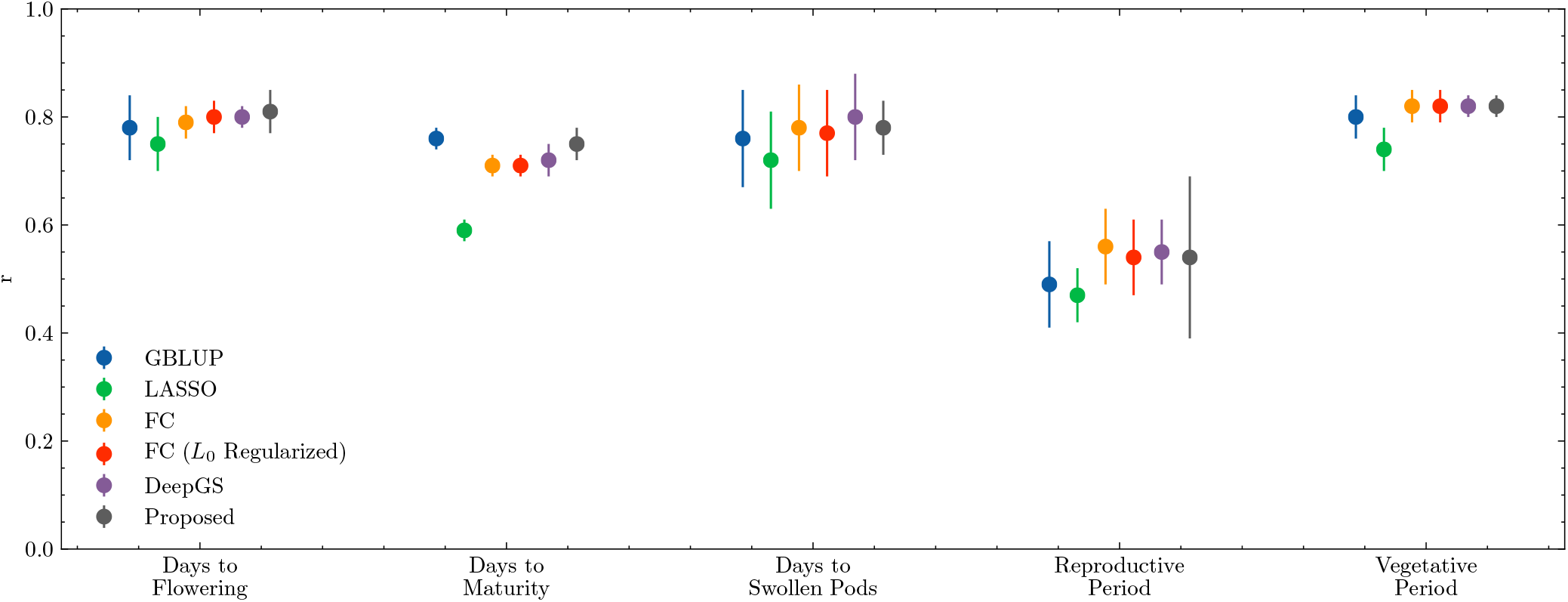
Genomic prediction results for the lentil diversity panel. Additional details are shown in Figure S1.

**Fig. 3.**
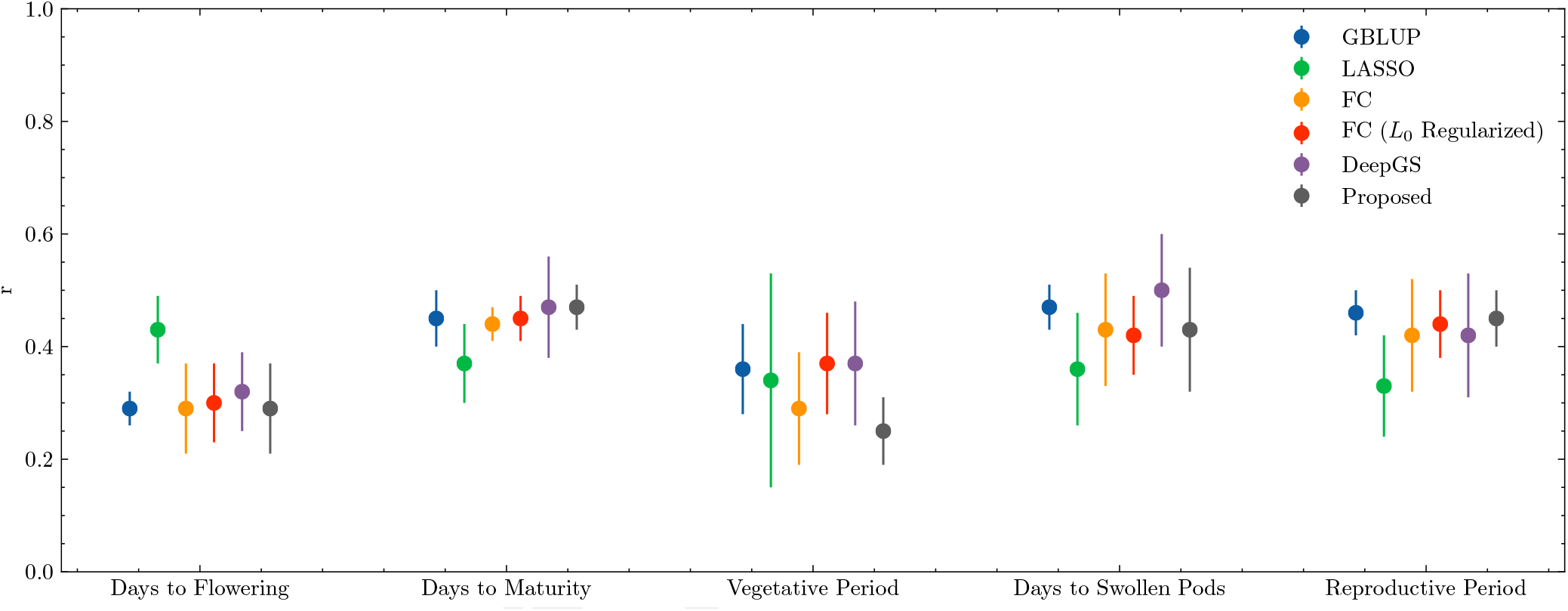
Genomic prediction results for lentil biparental populations. Additional details are shown in Figure S2.

**Fig. 4.**
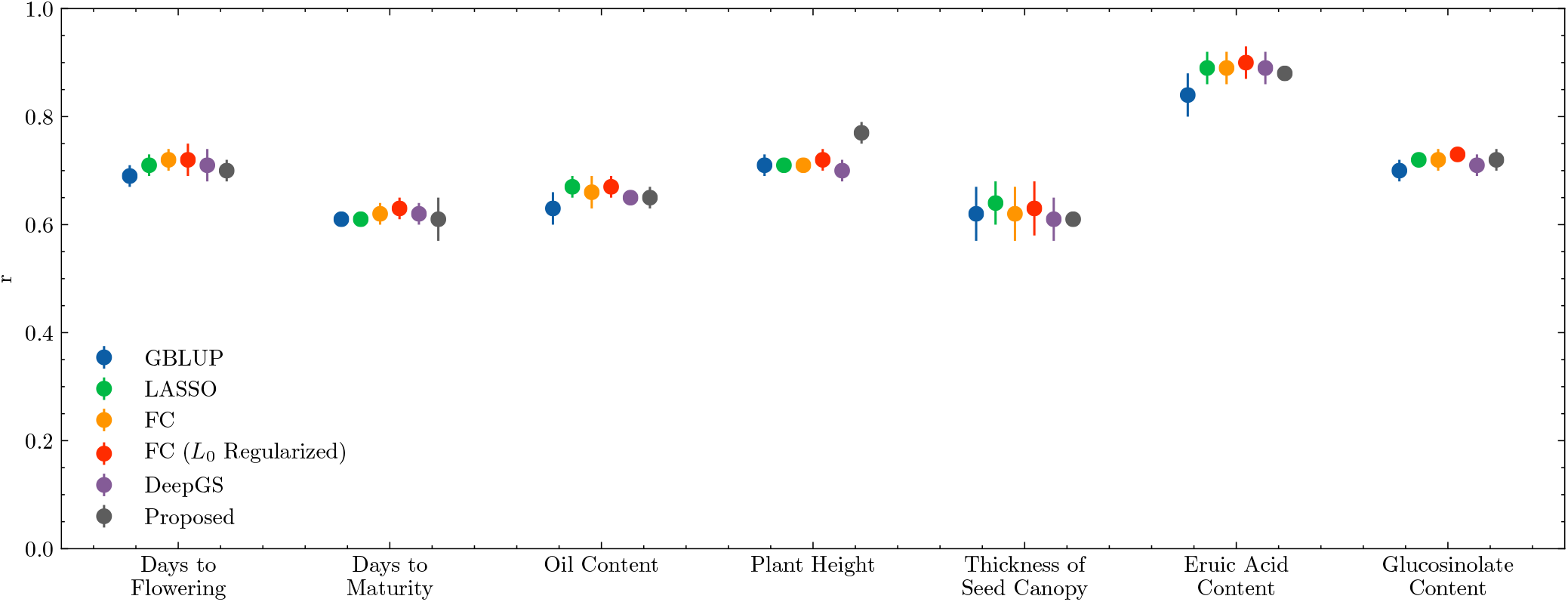
Genomic prediction results for *B. carinata*. Additional details are shown in Figure S3.

**Fig. 5.**
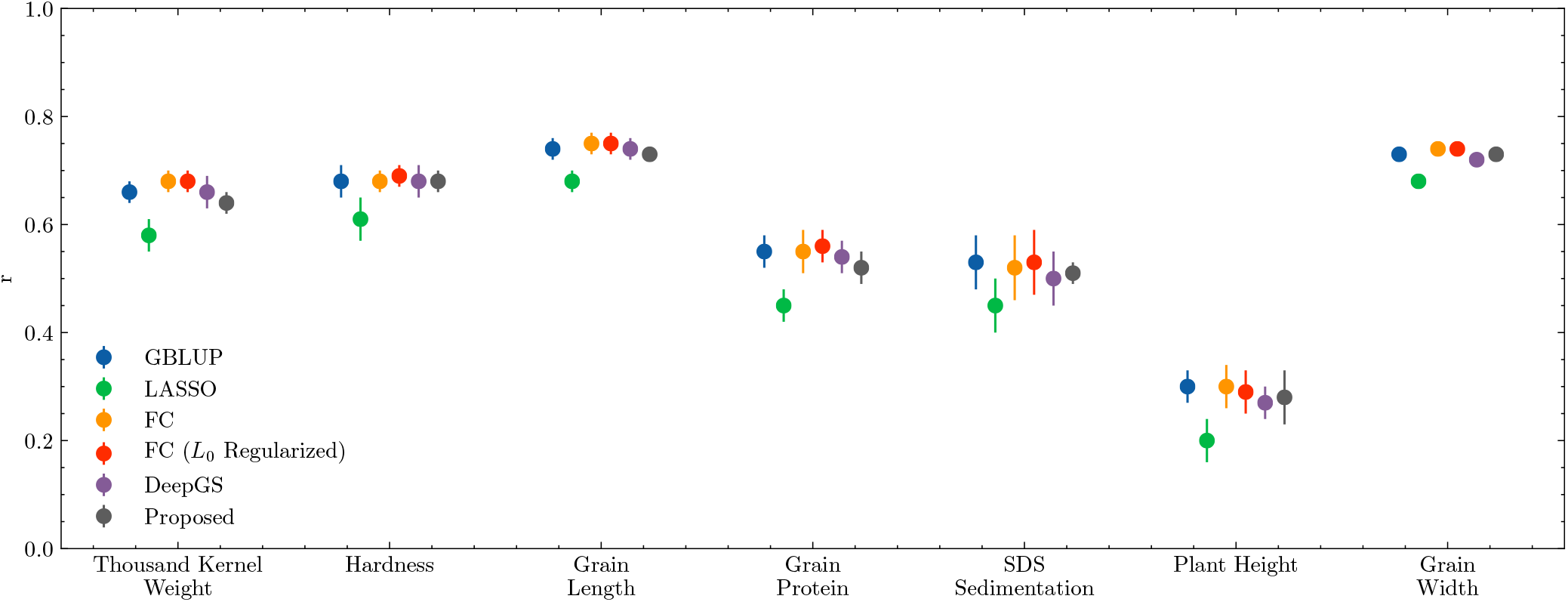
Genomic prediction results for wheat. Additional details are shown in Figure S4.

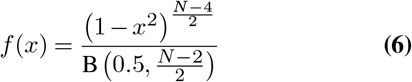

where B is the beta function and *N* is the sample size. Clearly, for small values of *N*, the expected variance in *r* values is high. We see this difference in variance between the biparental lentil dataset, which is the smallest in sample size, and the carinata and wheat datasets, which are significantly larger. In practice, we found *r* values to be volatile, varying considerably between different random splits of the data, and different random initializations. Even multiple runs with identical random seeds produce slightly different results for the machine learning models based on non-deterministic effects alone. For this reason, we decline to describe a method as superior when the difference in *r* is small, taking sample size into account. Due to the volatile and opaque nature of *r* as an estimate of predictive performance (for example, Figure 6), we also provide scatter plots of the results as supplementary figures.

**Fig. 6.**
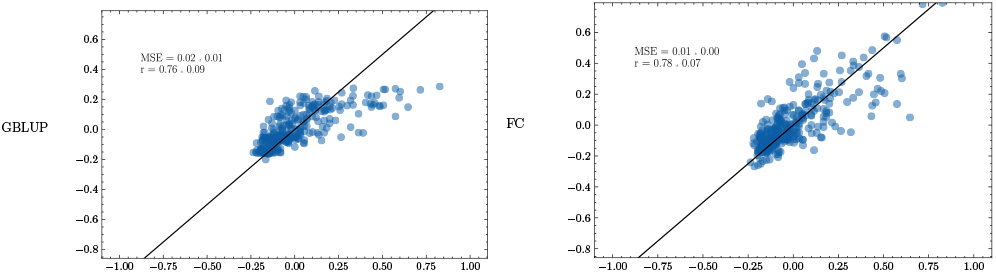
Example of scatter plots for the lentil diversity panel dataset using GBLUP (left) and FC (right). Although *r* is similar (0.76 and 0.78), the shape is different.

For all experiments, all six methods seem to provide similar predictive performance, with no method consistently outside the margin of error. The exception is LASSO, for which we observe a small decrease in performance on the LDP and wheat datasets relative to the other methods. LASSO also exhibits more volatile performance than the other methods across the experiments, likely due to the difficulties of training penalized linear models and their dependence on the strength of the regularization term. Among all the methods investigated in our evaluations, GBLUP continues to prove itself as a flexible choice when the underlying genetic structure of the trait is unknown. Although it is only capable of modeling linear additive effects with constant variance, GBLUP showed strong performance across species and traits. Despite this relatively good predictive performance, it remains the simplest method to implement, requires no tuning, and terminates in considerably less time than any of the machine learning methods.

Overall, our results largely mirror the findings elsewhere in the literature – for sufficiently large datasets, most methods offer comparable predictive performance regardless of differences in model capacity or complexity. Most importantly, the proposed method did not perform significantly worse than the standard DeepGS model on any dataset, even though it had no access to the marker data.

## Discussion

### On Shortcut Learning

It is reasonable to assume that, having the capacity to model complex epistatic effects between markers, deep neural networks would use this capability during training. However, deep neural networks are known to rely on features of the input which happen to correlate with the cost function, but which are not the features the network was intended to attend to. This phenomenon has been termed *shortcut learning*^1^ (23). For example, it has been shown that neural networks attend to color and texture more than the form of objects when performing image classification. Learning these simpler, higher-level features, or shortcuts, provides the network with a path of least resistance during optimization. The network does not need to learn a complicated representation of “cow” if the presence of cows is correlated with green grass in the image – looking for green pixels in the lower half of the image will suffice (23).

The shortcut learning principle may be true of genomic data as well. The network may find it unnecessary to seek out a small number of quantitative trait nucleotides (QTN), which are likely to be associated with a small and noisy gradient signal, when high performing individuals are likely to share large haplotype blocks in common by virtue of being genetically related. From the perspective of shortcut learning, it is not unlikely that the indifference of the network to the markers is due to the fact that it considers high-level genetic relatedness as a viable shortcut for predicting the phenotype.

### Limitations

One study reported that for simulated non-additive genetic architectures, neural networks (FC networks and CNNs) performed slightly better than GBLUP for a 100-QTN simulated trait, albeit slightly worse for a 1000-QTN simulated trait (9). This could be taken as evidence to suggest that there may be some nonlinear genetic architectures where deep neural networks are advantaged due to their ability to estimate nonlinear effects. However, it is worth noting that, for practically all real-world datasets (in this study, in (9), as well as elsewhere in the literature (6, 10–12)), this difference between neural networks and linear models is not observed. This discrepancy may be explained by differences in marker effect sizes between real and simulated traits, or how biologically plausible the architecture of the simulated trait was in practice. These issues could have an effect on the behavior of the network in terms of which features are salient for prediction. Finally, these small performance differences could also be accounted for by, as we’ve observed, volatility in Pearson’s *r*.

## Conclusion

Compared to linear models, deep neural networks have the ability to model complex nonlinear effects – however, this capability does not seem to benefit them in GP. In this paper, we show that one can obscure the marker data from the network, blinding it to which alleles are present in each individual – however, doing this does not seem to be at all deleterious to performance. Taken together, these two observations offer evidence that deep neural networks generally do not learn to model epistatic effects at all, but rather depend on estimating overall genetic relatedness. We suggest that that this is consistent with the behavior of neural networks in other domains, and is explained by a concept called shortcut learning.

This work reveals a clear direction for future work in deep learning for GP. The development of models which are capable of avoiding the learning of shortcuts in genomic data may unlock some of the potential of these powerful nonlinear models in genomic prediction.

## Data Availability

The data used in the wheat and lentil experiments are available from the sources referenced by their authors (21, 22). The carinata data is available on request.

## ACKNOWLEDGEMENTS

We thank Kirstin Bett for productive conversations about the lentil data. We also thank Mitchell Feldmann for his comments on the manuscript.

## Supplementary Figures

**Fig. S1.**
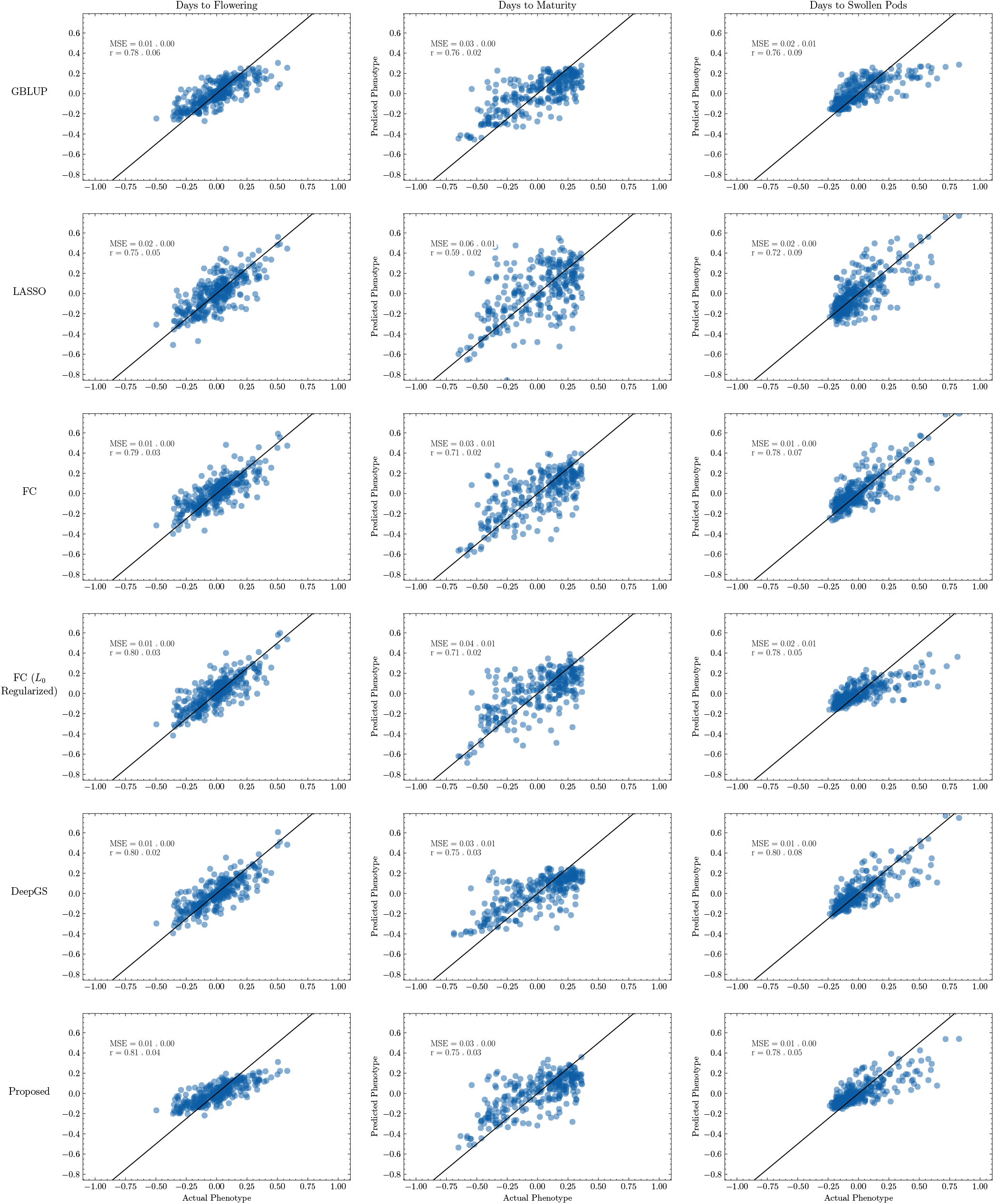
Detailed results for the first three traits in the lentil diversity panel dataset. The line *y = x* is shown in black.

**Fig. S2.**
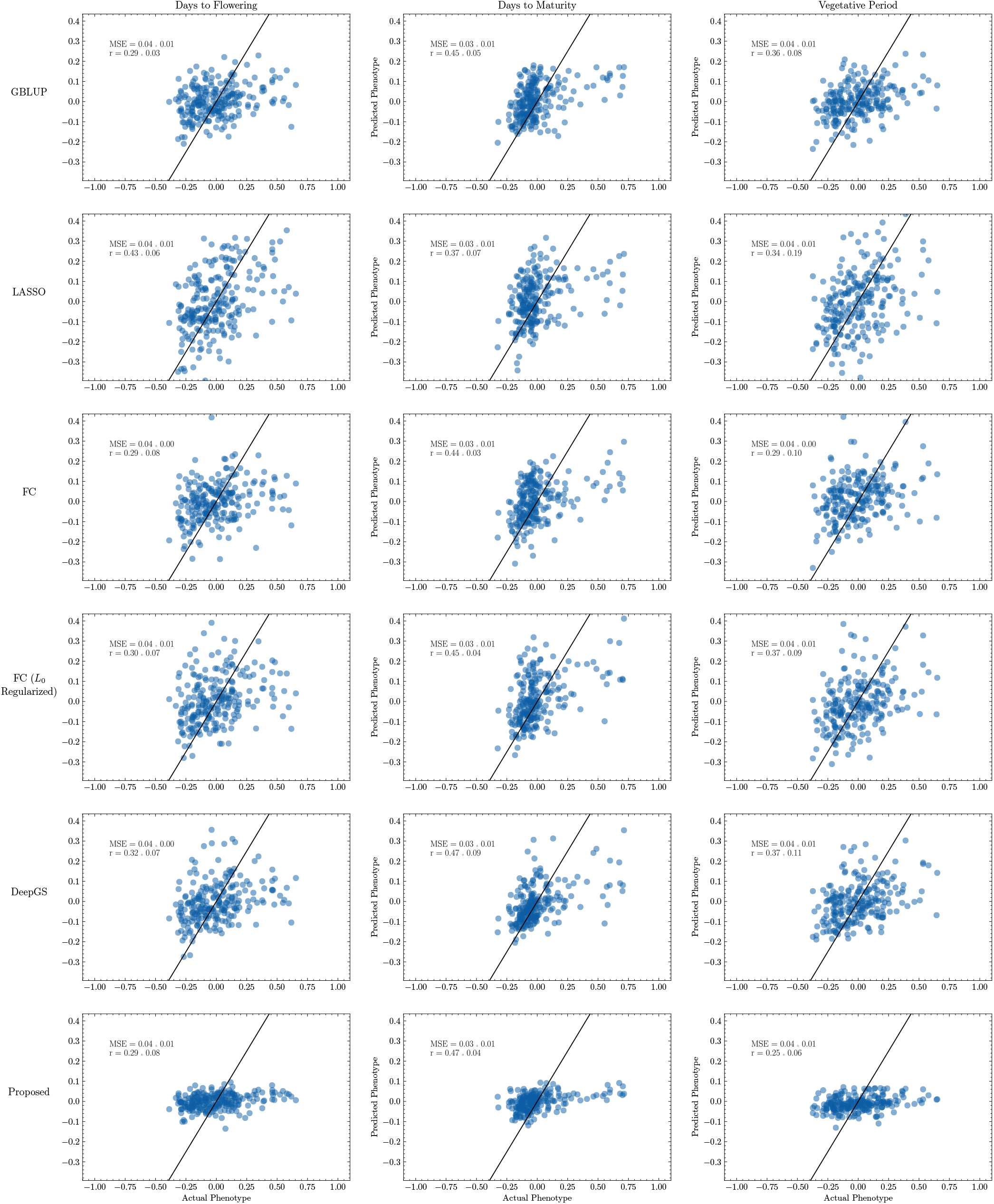
Detailed results for the first three traits in the lentil biparental dataset. The line *y* = *x* is shown in black.

**Fig. S3.**
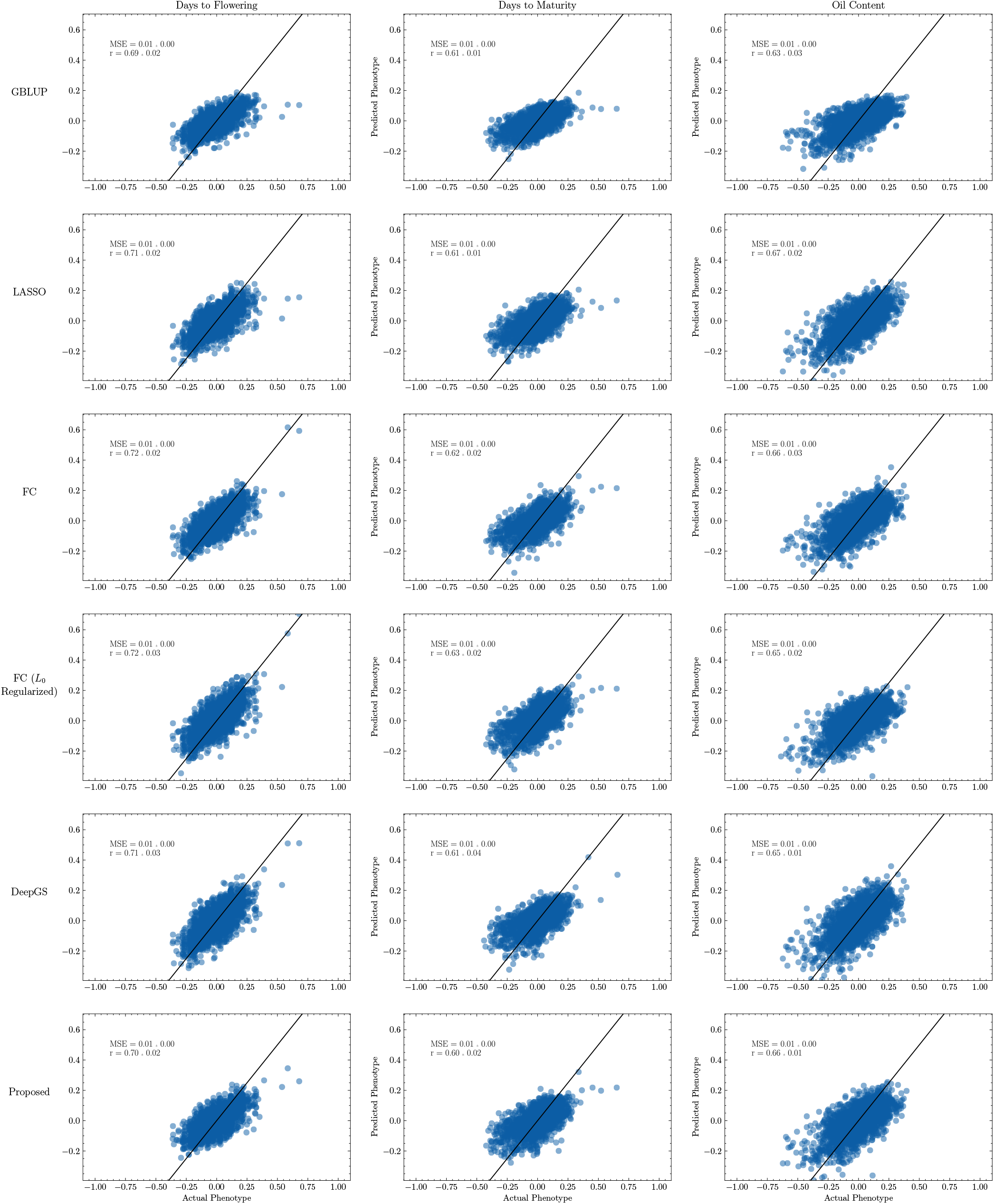
Detailed results for the first three traits in the carinata dataset. The line *y* = *x* is shown in black.

**Fig. S4.**
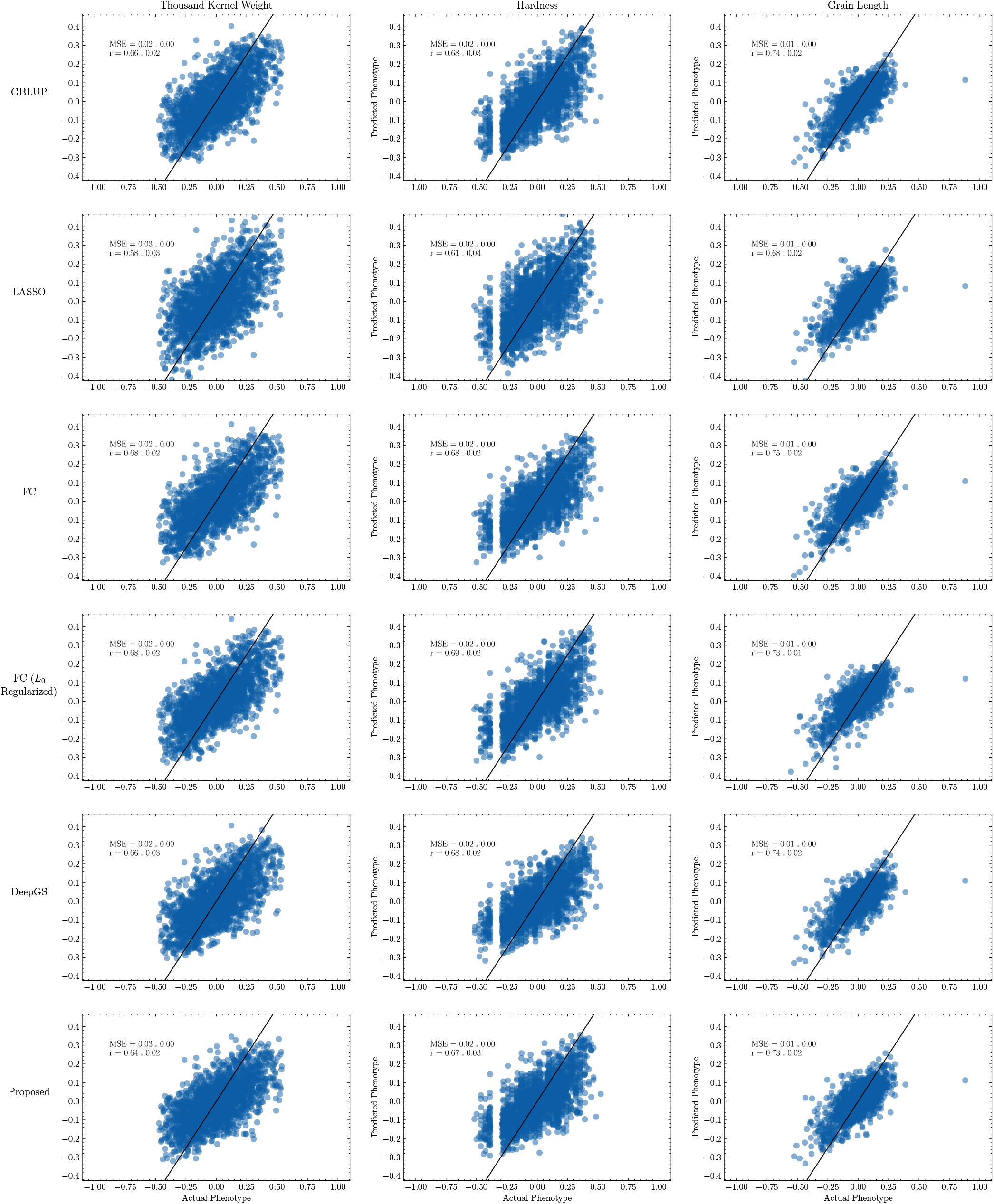
Detailed results for the first three traits in the wheat dataset. The line *y* = *x* is shown in black.

## Footnotes

1 This is distinct from the use of the term *shortcut* to refer to skip connections in some neural network architectures.

## Notes

### Competing Interest Statement

The authors have declared no competing interest.

